# Locus Coeruleus-Amygdala Circuit Disrupts Prefrontal Control to Impair Fear Extinction

**DOI:** 10.1101/2025.10.02.680131

**Authors:** Hugo Bayer, Annalise N. Binette, Samantha O. Sweck, Vitor A. L. Juliano, Samantha L. Plas, Lara M. Ferst, James E. Hassell, Stephen Maren

## Abstract

**Background:** Stress undermines extinction learning and hinders exposure-based clinical therapies for a variety of neuropsychiatric disorders. In both animals and humans, dysfunction in the ventromedial prefrontal cortex (vmPFC) contributes to stress-impaired extinction, but the neural circuit by which stress modulates vmPFC function is not known. We hypothesize that the locus coeruleus (norepinephrine system (LC-NE) undermines extinction learning by recruiting projections from the basolateral amygdala (BLA) to vmPFC.

**Methods:** We combined chemogenetics, calcium imaging, and fiber photometry to examine how the LC-NE system influences fear extinction, with special interest to LC→BLA projections. We infused viral vectors into the LC, BLA, and ventromedial prefrontal cortex (vmPFC) to express designer receptors (hM3Dq) or calcium indicators (GCaMP). The LC was globally or selectively (LC→BLA projections) stimulated, while vmPFC and BLA activity was monitored during different stages of memory processing. Intra-BLA propranolol infusions were used to block β-adrenergic receptors to test their role in LC-driven effects.

**Results:** We found that chemogenetic activation of the LC increased freezing behavior, suppressed vmPFC neuronal activity, and mimicked the effects of footshock. LC stimulation impaired both delayed and immediate extinction learning, while activation of the LC→BLA pathway alone was sufficient to drive the immediate extinction deficit. LC activation increased activity in BLA neurons projecting to vmPFC, and this effect, as well as vmPFC suppression, was prevented by β-adrenergic blockade with propranolol in the BLA. Overall, LC-driven NE release in the BLA disrupted vmPFC activity and dynamics, promoted a high-stress stated and impaired fear extinction.

**Conclusion:** This study demonstrates that stress and LC activation promote NE release in the BLA, which disrupts vmPFC activity and impairs fear extinction. These findings identify the LC–BLA–vmPFC circuit as a key pathway through which stress undermines extinction learning, highlighting BLA β-adrenergic receptors as potential therapeutic targets for stress-related disorders like PTSD.

## INTRODUCTION

After a threatening experience, it is crucial to organize defensive behaviors to mitigate harm. Once the threat passes, it is equally important to suppress fear responses that are no longer adaptive in the absence of danger. Suppression of conditioned fear involves extinction learning, which occurs when a stimulus that once predicted an aversive outcome no longer does so^1,2^. A vast body of evidence shows that suppressing learned fear is challenging: acquisition and expression of fear extinction are undermined by numerous factors, including stress^3^. For example, patients with post-traumatic stress disorder (PTSD), show resistance to extinction^4^. Indeed, extinction learning is thought to mediate exposure-based therapies used to treat PTSD and stress-induced impairments in extinction undermine the efficacy of these clinical interventions^5^. For example, extinction tends to be less effective when performed soon after an emotionally arousing event, including fear conditioning^6^. This “immediate extinction deficit” (IED) is mitigated by manipulations that reduce fear prior to extinction training^7^. The IED and other forms of stress-induced extinction impairments may contribute to the relapse of fear often seen in patients undergoing exposure therapy^8^.

The IED is associated with reductions in neuronal activity in the ventromedial prefrontal cortex (vmPFC), a brain region that is critical for extinction learning^9–12^. Animals exhibiting an IED show reduced levels of Fos express in the vmPFC^12^ and pharmacological activation of vmPFC neurons rescues the deficit^6^.Importantly, the IED is associated reduced spontaneous spike firing in the vmPFC^10^, mirroring other stress-induced disruptions of vmPFC anatomy and physiology^13^. Several studies have shown that propranolol, a ß-noradrenergic antagonist, mitigates stress-induced impairments by normalizing vmPFC firing to enable extinction learning^10,14–16^. Systemic propranolol not only mitigates stress-induced decreases in vmPFC activity but also reduces shock-induced increases in BLA spike firing that may contribute to the IED^14^. Interestingly, the effects of systemic propranolol on vmPFC activity appear to be mediated by the basolateral amygdala (BLA)^15^. This points to the BLA as a key mediator of stress-induced extinction deficits and as the link between peripheral stress responses, norepinephrine release, and prefrontal suppression.

One of the most important stress-sensitive systems in the central nervous system is the locus coeruleus-noradrenergic (LC-NE) system^17,18^. The LC is a small brainstem region that accounts for most of the forebrain norepinephrine^19^. The LC-NE system is important for the regulation of multiple behavioral domains, such as sleep and wakefulness, attention, anxiety, sensory arousal, and long-term memory^17,18,20–23^. Importantly, aberrant LC activity has been proposed to affect pathological anxiety and PTSD development^23,24^.

The recruitment of the LC-NE system is also important for fear conditioning. Footshock increases the release of NE in the basolateral amygdala^16,25,26^ and silencing LC⟶BLA projections during footshock impairs fear conditioning^27^. Additionally, it has been reported that intra-BLA injections of propranolol attenuate the IED^15^ and limit the induction of the IED after chemogenetic LC stimulation^14^.

On the other hand, NE release and adrenergic receptor activation in the medial prefrontal cortex (vmPFC) are necessary for extinction learning^28,29^, and silencing LC⟶vmPFC projections impairs extinction learning^27^. It has been proposed that the LC might switch from a “global broadcast mode” to a “discrete coding mode”. During intense threat (fear conditioning), the LC is broadly activated to mediate general arousal. In contrast, during early extinction (fear retrieval), its projections to the BLA are selectively engaged, whereas during late extinction its projections to the vmPFC are preferentially activated^27^. However, little is known about how stress-induced LC activity and increased norepinephrine release in the BLA affects other brain regions downstream of the BLA, like the vmPFC.

Here, we examined downstream effects of LC stimulation during extinction as well as in naïve animals. We used a chemogenetic approach to drive either global LC activation or targeted stimulation of LC⟶BLA projections to assess contributions of these projections to vmPFC activity and fear extinction learning using a combination of pharmacological and calcium imaging approaches. We found that chemogenetic stimulation of LC drove freezing behavior irrespective of learning, and both footshocks and LC stimulation led to similar changes in vmPFC activity. Global LC stimulation as well as stimulation of the LC⟶BLA projection led to impairments of delayed and immediate extinction learning and retrieval. Furthermore, LC stimulation combined with a weak foot-shock led to increased activity in vmPFC-projecting BLA neurons. Finally, we show that the reduced activity and changes in population dynamics in the vmPFC caused by LC stimulation are dependent on β-receptor activation in the BLA.

## MATERIALS AND METHODS

### Animals

In this study, we used 47 male and female Long-Evans rats obtained from Envigo (200–225 g on arrival). Rats were individually housed in cages within a temperature- and humidity-controlled vivarium and kept on a 14:10 light: dark cycle (lights on at 7 AM). Experiments were performed during the light phase of the cycle. Rats had ad libitum access to standard rodent chow and water. Before behavioral testing, rats were handled for 5 days (∼30 sec/rat/day). All experimental procedures were conducted in accordance with the US National Institutes of Health (NIH) Guide for the Care and Use of Laboratory Animals and were approved by the Texas A&M University Institutional Animals Care and Use Committee (IACUC).

### Surgery

Rats were anesthetized with isoflurane (5% for induction and 1–2% for maintenance) and secured in a stereotaxic apparatus (Kopf Instruments, Tujunga, CA). Animals received a 10 mg/ml/kg injection of Rimadyl (Zoetis), the incision area on the scalp was shaved, povidone-iodine pads were used for antisepsis, ophthalmic ointment was applied to the eyes, and lidocaine was injected subcutaneously beneath the scalp. An incision was made using a scalpel blade and the scalp was retracted using hemostats. The precise procedures for each surgery involved in the study are described below.

For virus infusions, a 27G injector needle was connected to a PE10 tubing (Braintree Scientific) that was connected to a Hamilton syringe in the other end, and the Hamilton syringe was mounted in a Legato 101 infusion pump (KD Scientific) that was used to determine the volume and rate of infusion. For LC infusions, the head was tilted down at a 15o angle so that the bregma was 2 mm below the lambda in the DV axis, and bilateral infusions of AAV9-PRSx8-hM3Dq-HA (1.214e14 GC/mL, University of Pennsylvania Vector Core) were made using the following coordinates [mm relative to lambda]: AP = -3.8, ML = ± 1.4, DV = -7.0, -6.5, and -6.0. Three virus infusions of 0.5 µl each were used to facilitate targeting of the LC, and do not lead to off-target expression since the virus carries the PRSx8 promoter, which restrict expression to NE+ cells since it is a dopamine-β-hydroxylase promoter^58^. Animals were given at least 2 weeks to allow for viral expression before the beginning of behavioral experiments. For BLA infusions, bregma and lambda were leveled, and infusions were made using the following coordinates [mm relative to bregma]: AP = -2.85, ML = ± 4.85, DV = −8.7.

In different experiments, the BLA was bilaterally infused with CAV2-PRSx8-hM3Dq-mCherry (0.5 µl/side; 3e12 GC/mL, Plateforme de Vectorologie de Montpellier), CAV2-mCherry (0.5 µl/side; 3e12 GC/mL, Plateforme de Vectorologie de Montpellier), AAV9-CAG-FLEX-jGCaMP8f-WPRE (0.75 µl/side; 5e12 GC/mL, Addgene plasmid #162380), and AAV9-AAV-hSyn-GRAB-NE1m (0.5 µl/side; 1e13 GC/mL, Addgene plasmid #123308). For the vmPFC infusions of CAV2-Cre (0.5 µl/side; 5e12 GC/mL, Plateforme de Vectorologie de Montpellier), we used the following coordinates [mm relative to bregma for AP and ML, and skull surface for DV with a 30o angle]: AP = +2.65, ML = ± 3.2, DV = -6.5. For calcium imaging experiments the vmPFC was unilaterally infused with AAV1-CaMKIIa-GCaMP6m-WPRE-SV40 (1µL; 3.8 x 1012 GC/mL, Inscopix).

For BLA cannula implantation, one stainless steel guide cannulae (26 G, 9-mm from base of the pedestal; RWD) was implanted unilaterally in the BLA (same hemisphere in which GRIN lenses were implanted in the vmPFC) using the following coordinates [mm relative to bregma for AP and ML, and relative to skull surface for DV]: AP = -3.5, ML = ± 2.85, DV = −8.55. For GRIN lens implantation in the vmPFC, a burr hole was drilled using the following coordinates [relative to bregma]: AP = 2.7; ML = ±0.5. Then, a 22G needle was lowered at the rate of 0.2 mm per minute until it reached the -4.4 in the DV axis relative to dura and slowly pulled up after that. The 0.6 × 7.3 mm GRIN lens was then placed over the craniotomy and lowered to touch dura and then lowered at the rate of 0.1 mm per minute until it reached -4.4 DV relative to dura. For GRIN lens and cannula implantation surgeries, 3 screws were attached to the skull to help secure the headcaps. After surgeries, topical antibiotics (Triple Antibiotic Plus, G&W Laboratories) were applied to the skin surrounding the headcap.

### Drugs and infusion procedures

For chemogenetic stimulation of the LC, rats received intraperitoneal (i.p.) injections of clozapine-N-oxide (CNO) dissolved in DMSO (2.5% of total volume) at the dose of 3 mg/kg and the volume of 1 mL/kg. 0.5% DMSO saline was used as vehicle for control animals. For intra-BLA propranolol infusions, animals were transported to a dedicated infusion room and stainless-steel injectors (33 G) connected to Hamilton syringes mounted on infusion pumps were inserted into the guide cannulae. Propranolol was dissolved in saline to reach the concentration of 5 µg/µL, and 0.5 µL were injected into the BLA at the rate of 0.25 µL/min, and after the injections, cannulas injectors were left in place for an additional minute. In control animals, saline served as the vehicle.

### Behavioral apparatus and procedures

Fear conditioning was conducted in 16 standard rodent conditioning chambers (30 × 24 × 21 cm; Med Associates, St Albans, VT), housed inside sound-attenuating cabinets. The chambers had two aluminum sidewalls, a Plexiglas rear, ceiling, and door, and a grid floor. The grid floor consisted of 19 stainless steel bars that were connected to a shock source, and a solid-state grid scrambler for foot shock delivery (Med Associates). Each conditioning chamber was equipped with a loudspeaker for delivering auditory stimuli, a ventilation fan, and a light. Locomotor activity was transduced into an electrical signal by a load cell under the floor of the chamber to automatically measure freezing. Rats underwent auditory fear conditioning in context A (chamber lights on, fans off, cabinet doors closed, 3% acetic acid, white transport boxes), which consisted of five tone-footshock pairings after a 3 min baseline period. The tone conditioned stimulus (CS) was 10 s, 80 dB, 2 kHz; the shock unconditioned stimulus (US) was 1 mA, 2 s; the intertrial interval (ITI) was 70 s and animals remained in the chamber for an additional minute after the last pairing. Context exposure consisted of a 10-min stimulus-free period in the extinction context (context B; chamber lights off, fans on, cabinet doors open, 70% ethanol, black floors covering the grid floor, and black transport boxes with bedding). Extinction and extinction retrieval consisted of a 3-min stimulus-free baseline period followed by 45 CS-alone presentations (ITI = 40 s) followed by a 3 min post-trial period. Immediate and delayed extinction were identical except for the fact that immediate extinction was performed 15 min after conditioning, while delayed extinction was performed one or two days later.

For two experiments, animals were conditioned using a weak conditioning protocol. That procedure was similar to the regular conditioning protocol described above, except that during conditioning, a single CS-US association was made, the CS was shorter (2 s) and the footshock was shorter and weaker (0.5 mA, 0.5 s), and the CS consisted of 80 dB white noise instead of a 2 kHz pure tone. This procedure was previously used in our laboratory and led to low levels of conditioned fear and was not enough to produce the immediate extinction deficit14.

For some of the calcium imaging experiments, we recorded vmPFC activity under a variety of conditions, including naïve animals, unsignaled shocks, as well as during fear extinction. The precise moment of recordings is described for each experiment. Briefly, in experiments involving measures of vmPFC spontaneous activity after fear conditioning, animals were shocked and vmPFC activity was recorded 15 minutes afterwards, so that vmPFC measures correspond to a point in time animals would be undergoing immediate extinction. In experiments involving vmPFC recordings after LC chemogenetic stimulation, animals were injected with CNO, and we waited 15 min to start recordings. For experiments involving calcium imaging during extinction and retrieval, animals were extinguished in the calcium imaging chamber, and we recorded throughout the session. For all experiments, animals had a baseline recording session that lasted 5 minutes, which was used to verify that calcium signals are present, and before extinction and retrieval sessions, a 5-minute stimulus-free period served as a baseline for vmPFC activity measures.

For the experiment involving GCaMP fiber photometry, animals underwent the weak conditioning protocol described above. CNO was injected 10 minutes before the animals were placed in the conditioning chamber. After a 3 min baseline period, animals were fear conditioned using the weak conditioning protocol, and after fear conditioning, the animals remained in the chamber for 15 minutes before extinction started. All of that happened in the same chamber so that we could record changes in calcium transients throughout conditioning, interval and extinction.

### Calcium imaging and fiber photometry

Calcium imaging data were collected with an Inscopix imaging system (nVoke) and Inscopix Data Acquisition Software. Calcium transients were collected at a 5-10Hz rate. Optimal light settings (i.e. power, gain, and focus) were determined for each animal, and these settings were kept consistent across sessions. A TTL signal was used to synchronize calcium imaging and behavioral recordings. Data were analyzed with Inscopix Data Processing Software. Preprocessing algorithms were used to remove artifacts (early frames trimmed and defective pixels fixed) and reduce the movie size (cropped and spatially downsampled ×4). A spatial filter algorithm was used to remove low and high spatial frequency content. A motion correction algorithm was used to reduce frame-to-frame motion; the mean image was used as the global reference frame origin. The constrained non-negative matrix factorization (CNMF-E) algorithm was used to identify the spatial location of cells and their associated calcium traces. The CNMF-E extracted calcium traces were divided by their respective estimated noise. Lastly, traces were deconvolved using the OASIS algorithm to estimate neural spikes (spike SNR threshold: 6). Identified cells were manually inspected and false positive cells were excluded from analyses.

A fiber photometry system (FP3002; Neurophotometrics; San Diego, CA) was used to measure GCaMP8f and GRAB-NE1m signals in the BLA. LEDs (470 nm and 410 nm) were used as photoexcitation for the GCaMP8f and isosbestic signal, respectively. The light intensities of the source LEDs were calibrated to obtain ∼ 50 μW power at the tips of optical fibers. Sampling rate was set at 40 Hz for 410 nm and 470 nm LEDs. Raw fiber photometry data were then transferred to ΔF/F values normalized by the isosbestic signal by using pMAT, an open-source analysis software^59^. Calcium traces were then normalized based on the baseline period for the analysis shown in Figure 5D.

### Freezing onset correlation with vmPFC activity

For behavioral correlations with calcium imaging data, raw movement data obtained from the load cell was aligned to calcium transients and calcium transients between freezing and non-freezing periods were compared to determine whether there were differences between behavioral epochs. For the freezing onset analysis, only freezing bouts that had at least 3 seconds of non-freezing before the onset and lasted at least 5 seconds were included, and the normalized calcium transients were used to determine pre-onset slopes. The pre-onset slope was calculated based on the average of the 1st derivative of the calcium transient between -0.8 s and -0.2 s before freezing onset^60^. For the comparison with pre-onset slopes, we randomly sampled 100 period of similar length (8 seconds) from each animal and compared the slopes of the same period during those periods to the pre-onset slopes.

### Peri-event time histograms and neural trajectories

For peri-event time histograms (PETHs), calcium imaging data corresponding to early extinction, late extinction, and retrieval trials was aligned to CS onset. For each CS, data was z-scored using the 2 seconds before CS onset as baseline. For early and late extinction, the z-scored data of the first and last 5 CS presentations of the extinction session were averaged, and for the retrieval PETHs, the first 5 CS presentations of retrieval were averaged. The comparison between VEH and CNO PETHs was made by calculating the PETH area under curve (AUC) for each animal and comparing AUC group averages.

For analyzing neural CS neural trajectories, neurons from all animals, conditions and trials (early and late extinction and retrieval trials) were pooled and a single principal component analysis (PCA) was performed in the entire dataset so that the same principal components could be used across sessions and animals^61^. The total number of neurons included in the analysis was matched to the subject with the smallest number of neurons, and the subjects with higher numbers were randomly down sampled to match that number. The neural trajectories were created using the averaged calcium event number for 200ms windows starting at 2 seconds before CS onset and ending at 5 seconds after CS onset. The trajectories were plotted in PCA space using the PCA coordinates for each time-point. For each trajectory, the length was calculated as the sum of Euclidean distances between adjacent time-points. To statistically compare trajectories between the VEH-treated and the CNO-treated groups, we used the leave-one-out method, that consists of leaving all neurons from each animal per iteration. For the trajectory length calculations, we used the number of principal components (PCs) that explained 95% of variance in the data, but for visualization we used the first three PCs.

### Population vectors and hierarchical clustering

To produce a population vector for each condition, neuronal firing rates were binarized such that each neuron was classified as active in a condition if it was detected during that condition. The resulting activity matrix was standardized and a PCA was performed in the entire remaining dataset. Then, neurons were projected into a two-dimensional PCA space, where their condition-specific firing was represented using pie chart symbols that were color coded to represent the conditions in which each neuron fired. For each condition, the mean PCA coordinates of active neurons were calculated, and for each condition one vector was plotted from the origin to illustrate condition centroids. To quantify differences relative to a reference condition, we calculated the mean PCA position of neurons active in the VEH x VEH group as a reference point. We then computed Euclidean distances from this reference for each condition to determine the average distance from each condition centroid to that of the VEH x VEH condition.

To create a Venn diagram, we determined the overlap of active neurons across experimental conditions by first converting firing rates into binary values, with neurons classified as active if they fired at least once while the animal was in each experimental condition. This was also used to create a binary matrix with neurons as rows and conditions as columns. Pairwise Jaccard similarity coefficients were then calculated between conditions to quantify the proportion of shared active neurons relative to the union of active neurons. The resulting similarity matrix was visualized using a heatmap, and hierarchical clustering (Ward’s method) was applied to the corresponding distance matrix (1 – similarity) to generate a dendrogram of condition clusters.

### Network analysis

For analyzing network properties among groups, a cooccurrence matrix was created for each condition for each animal62. For that, calcium traces from all cells detected in that session were aligned and calcium events were extracted using the same criterion as described above (SNR > 6). Neuronal data was then converted to a binary dataset that had calcium events detected in 200 ms bins (so that each neuron was assigned 0 or 1). Then, a symmetric co-occurrence matrix was created and neurons that fired within one second of each other were assigned a 1 in the co-occurrence matrix, otherwise they had a 0 for that window (Figure 4G). Data from all co-occurrence matrixes created were combined into a single matrix for the entire session, and that combined matrix was used to create the networks. Networks were created using the Python library NetworkX. In the network, each node is a neuron and an edge is created whenever two neurons had a 1 value in the combined co-occurrence matrix. The position of neurons in the network does not correspond to anatomical positions or coordinates in the calcium imaging video. For comparing networks among themselves, we used the Jaccard index which compares the edges of two networks and is calculated by dividing the number of common edges (intersection) by the total number of unique edges (union) to quantify network similarity between conditions. Additionally, the clustering coefficient for each network was calculated, which measures the proportion of a node’s neighbors that are connected to each other, which corresponds to the network’s tendency to form closely connected groups.

### Histological procedures

Rats were overdosed with sodium pentobarbital (100 mg/kg) and perfused transcardially with 0.1 M PBS followed by 10% formalin. Brains were extracted from the skull and post-fixed in a 10% formalin solution for 24 h followed by a 30% sucrose solution where they remained for a minimum of 48 h. After the brains were fixed, coronal sections (30μm thickness) were made on a cryostat (−20°C), mounted on subbed microscope slides, visualized in a fluorescent microscope (Zeiss Axioscope 5). Only animals with confirmed viral expression and implant placement were included in the analyses.

## RESULTS

### Chemogenetic stimulation of LC increases freezing and suppresses vmPFC activity in naïve rats

First, we characterized the effects of global chemogenetic LC stimulation on freezing behavior and vmPFC activity using naïve animals. For this purpose, 14 rats were implanted with GRIN lenses in the vmPFC and received infusions of AAV1-CaMKIIa-GCaMP6m-WPRE-SV40 in the vmPFC and AAV9-PRSx8-hM3Dq-HA in the LC (Figure 1A). Four weeks after surgery, animals were habituated to the miniscope tether for 3 consecutive days. After this habituation period, they underwent two recording sessions (Figure 1B). First, they were placed in a novel context for a 10-min recording session—this served as a baseline recording. After this, they were removed from the chamber, injected (i.p.) with vehicle (VEH), and immediately returned to the chamber for a 15-min post-injection recording session. This procedure was repeated on day two with CNO injections instead of VEH.

**Figure 1.**
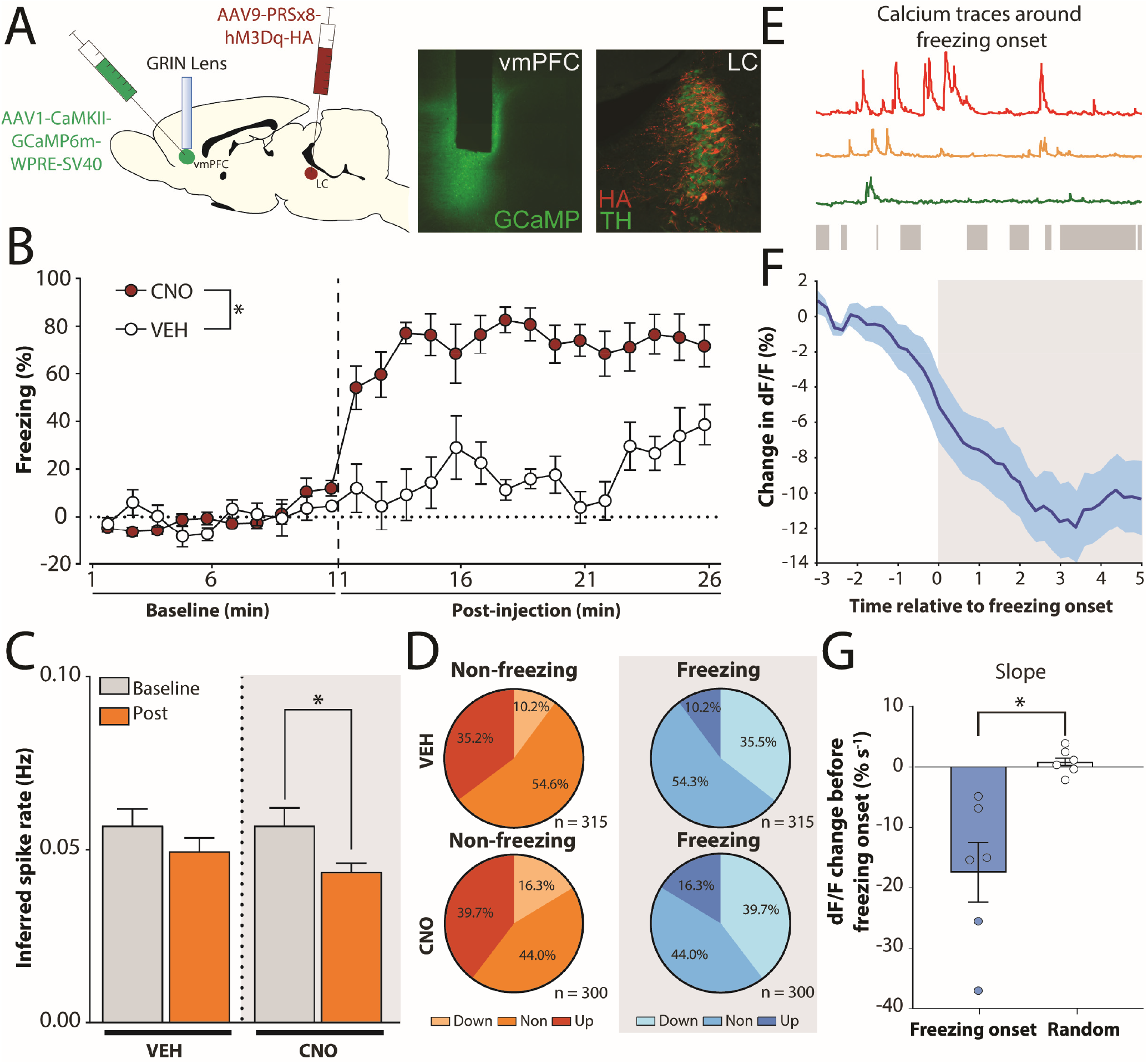
Locus coeruleus chemogenetic stimulation drives freezing behavior and suppresses prefrontal activity. (A) Surgery schematics and representative histology of locus coeruleus and medial prefrontal cortex. A DREADD-carrying vector with specificity to NE+ neurons was injected in the LC, a GCaMP vector was injected in the vmPFC, and a GRIN lens was implanted on the vmPFC of (n = 7; 4 female and 3 male). (B) Naïve animals present increased levels of freezing in a neutral context after CNO injections. (C) Inferred vmPFC spike rate is decreased after CNO, but not VEH injections (n = 260 cells for VEH and 257 cells for CNO). (D) Change in activity in vmPFC cells during non-freezing and freezing states after VEH or CNO injections. (E) For freezing onset analysis, activity from individual vmPFC cells was averaged and aligned with freezing bouts (gray bars). (F) When population calcium traces were aligned with freezing onset, there is a reduction in vmPFC activity around freezing onset, which is sustained through the freezing bout. (G) The slope of the calcium trace curve between -1.0 s and -0.2 s relative to freezing onset is significantly lower than slopes that were randomly sampled regardless of freezing. BL = baseline, CNO = clozapine-N-oxide; LC = locus coeruleus; vmPFC = ventromedial prefrontal cortex, * = p < 0.05, line and bar plots represent mean ± s.e.m.s.

CNO injections led to a robust increase in freezing behavior during the post-injection period relative to VEH injections (Figure 1B). Although no differences were observed during the baseline period (*F*_1,6_ = 0.8891, *p* = 0.38), a two-way ANOVA performed on the post-injection freezing levels revealed a main effect of CNO (*F*_1, 6_ = 85.82, *p* < 0.01), but no significant interactions(*F*_14, 84_ = 14.84, *p* = 0.09). CNO injections also reduced neuronal activity in IL: animals administered CNO showed a significant reduction in inferred spike rates in the post-injection period (Figure 1C). This was confirmed by *t*-tests that showed no significant difference in spike rate in the VEH condition (*t* = 1.82, *p* = 0.08), but a significant decrease in spike rate in the CNO condition (*t* = 3.47, *p* < 0.01). Additionally, we performed a permutation analysis by shuffling firing rates from baseline and post-injection periods 1000 times to generate a null distribution and used a *Z*-test for both the VEH and the CNO conditions (Figure S1). With that analysis, we confirmed that the post-injection firing rate for the VEH condition was not significantly lower than chance (*Z* = - 1.18, *p* = 0.24), while the firing rate for the CNO condition was significantly lower when compared to the null hypothesis (*Z* = -2.18, *p* = 0.03).

We next analyzed inferred spike firing rate in individual neurons recorded during the session to determine whether vmPFC activity is correlated with freezing behavior. Neurons were classified as up-, down-, and non-modulated cells based on the effect of VEH or CNO injection on their firing rates. Figure 1D shows the proportion of these cell classes observed during either freezing or non-freezing periods. When freezing and non-freezing periods are isolated, there is a reduction in vmPFC activity when animals are freezing, regardless of treatment (*F*_1, 613_ = 61.18, *p* < 0.01). These results show that the high stress state induced by LC chemogenetic stimulation is correlated with reduced vmPFC activity.

That observation raises the question of whether this reduction is a consequence of freezing behavior or whether it reflects the neural process that leads to the initiation of freezing. To address this question, we averaged calcium traces and aligned those with freezing bouts for each animal from both VEH and CNO conditions (Figure 1E, F). We then aligned the average dF/F trace with all “valid” freezing bouts for each animal (see methods), we observed a marked reduction in the average calcium signal during freezing events, and analyzed the period between -1.0 and -0.2 s relative to freezing onset to determine whether the reduction in calcium traces preceded freezing initiation or was caused by it (Figure 1F). The slope of pre-freezing onset periods was averaged to derive a single value per animal, and 100 calcium traces of the same length were randomly sampled for each animal to determine whether the average slope before freezing onset was different from chance. As seen in Figure 1 G, a *t*-test revealed a significant difference between freezing onset slopes compared to the randomly sampled (*t* = 3.68, *p* < 0.01). Together, our results show that LC chemogenetic stimulation in naïve animals leads to an increase in freezing behavior and this is correlated with reduced vmPFC activity. Interestingly, reduced vmPFC activity is more pronounced before and during freezing bouts.

### Chemogenetic stimulation of LC and fear conditioning lead to similar changes in vmPFC activity

Having discovered that chemogenetic LC stimulation induces a high fear state, we compared vmPFC activity in response to LC stimulation and footshock. To address this question, the same rats reported in Figure 1 were submitted to a 3-day experimental procedure (Figure 2 A, B). This consisted of 1) a VEH injection followed by exposure to the conditioning context and CS delivery without any footshocks on Day 1, 2) a CNO injection followed by exposure to the conditioning context and CS delivery without footshocks on Day 2, and 3) a VEH injection followed by a standard fear conditioning procedure on Day 3. Calcium signals were obtained during a 5-min baseline session before trials started and 15 minutes after injections (about 5 minutes after the last trial).

**Figure 2.**
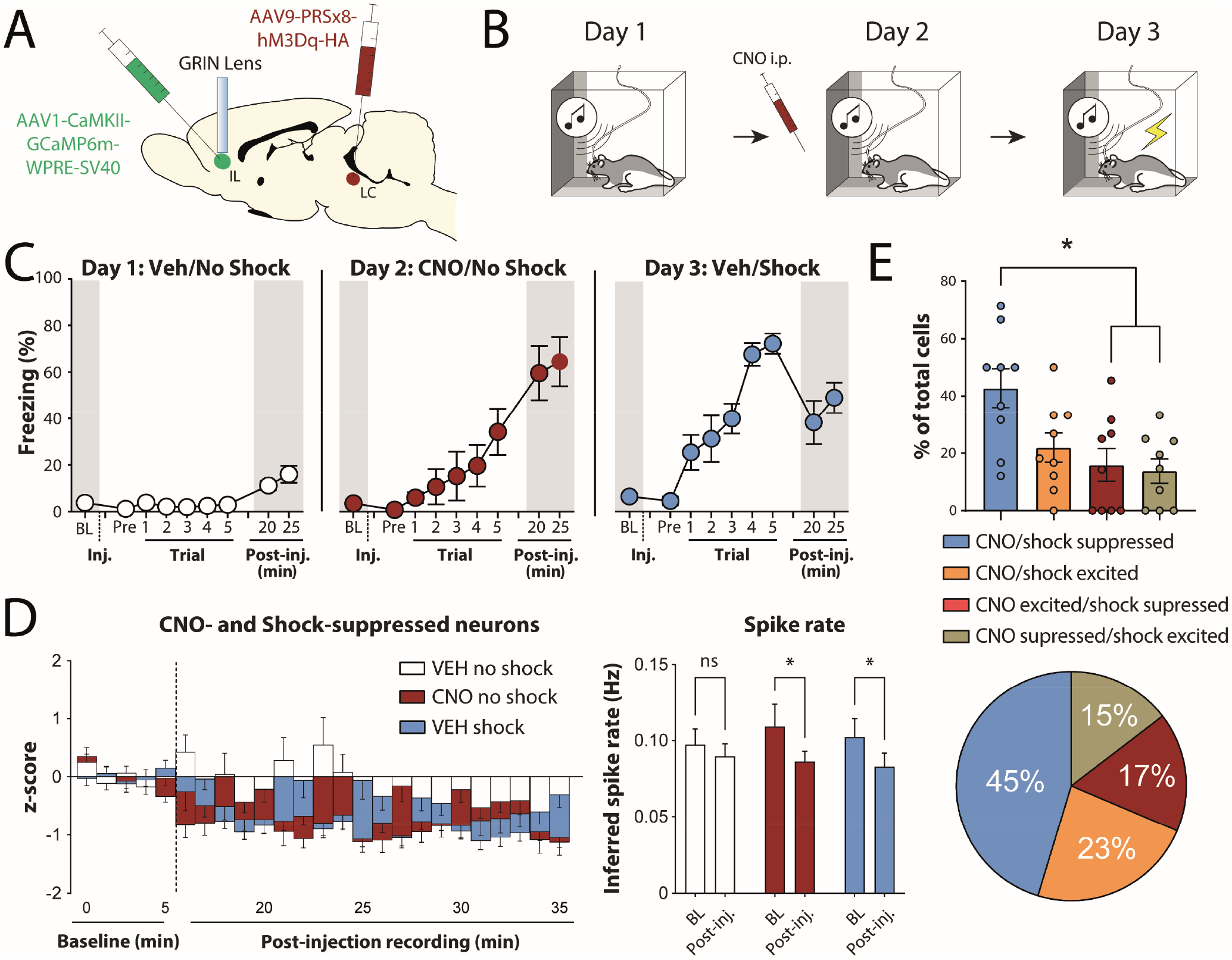
Locus coeruleus chemogenetic stimulation and shock increase freezing behavior and suppress vmPFC activity. (A) Surgery schematic. A DREADD-carrying vector with specificity to NE+ neurons was injected in the LC, a GCaMP vector was injected in the vmPFC, and a GRIN lens was implanted on the vmPFC (*n* = 9; 5 female and 4 male). (B) Behavioral procedure. On the first day, animals initially went into the conditioning chamber for a 5-min baseline (BL) period, injected with VEH, and returned to the chamber, where tones were played but no shocks were given. On the second day, the procedure was the same, but they were injected with CNO instead of VEH. On the third day, they once again received VEH injections, but underwent a conditioning session involving tones and shocks. (C) Freezing behavior across days. Animals presented very little freezing on the first day, but both CNO and shocks were able to drive considerable amounts of freezing during days 2 and 3. (D) *Left panel*, normalized mPFC calcium activity across baseline and post-injection periods; *right panel*, both CNO and shock reduced inferred spike rate in the vmPFC during the post-injection period, while activity was not significantly different in the VEH no shock condition (*n* = 168 cells). (E) Proportion of cells that are excited and/or suppressed by CNO and shock. There was a significantly higher proportion of neurons suppressed by both CNO and Shock compared with the other classes of neurons based on CNO and Shock response. BL = baseline, CNO = clozapine-*N*-oxide, LC = locus coeruleus, ns = not significant, VEH = vehicle, vmPFC = ventromedial prefrontal cortex, * = *p* < 0.05, line and bar plots represent mean ± s.e.m.s.

As shown in Figure 2C, both systemic CNO administration and footshocks led to increased freezing across the CS trials. This was confirmed by three separate one-way ANOVAs (CNO: *F*_6, 56_ = 2.62, *p* = 0.03; Shock: *F*_6, 56_ = 20.08, *p* < 0.01; VEH/No Shock: *F*_6, 56_ = 0.63, *p* = 0.70). When the freezing levels during the period after trials were analyzed with a one-way ANOVA, we observed a high freezing level in both the CNO and Shock conditions in comparison with the VEH/No Shock condition (*F*_2, 34_ = 25.19, *p* < 0.01), and a Tukey’s *post hoc* test revealed that both CNO and Shock conditions had higher freezing levels (*p*’s < 0.01). We then compared the inferred spike rate after trials across conditions, comparing the post-injection activity with the baseline for each condition (Figure 2D). The VEH/No Shock condition did not have significant differences in spike rate when baseline and post-injection were compared (*t* = 1.11 *p* = 0.27). However, in both the CNO and Shock conditions, we observed a decreased spike rate when the post-injection period was compared to the baseline (*t*’s = > 2.80, *p*’s < 0.01). Collectively, these data suggest that both chemogenetic stimulation of the LC and footshocks lead to a high fear state that is correlated with decreased vmPFC activity.

We then analyzed the individual neuronal activity across days to determine the shifts in neuronal activity in each condition. For this, we analyzed the difference in individual *z*-scored neuronal activity between baseline and post-injection periods. As shown in Figure 2E, the highest proportion of cells that showed common changes in activity to the drug treatment and behavioral manipulation were those cells that showed decreases in activity to both CNO and footshock. This was confirmed by a one-way ANOVA showing a significant main effect of group (*F* _*3,32*_ = 5.72, *p* <0.01). Additionally, a Tukey’s *post hoc* test revealed that the CNO/shock suppressed cells were more greatly represented in the population than cells suppressed in one condition and excited in the other (*p*’s < 0.01) and a trend towards significance was observed when CNO/shock suppressed cells were compared to the proportion of CNO/shock excited cells (*p* = 0.056). These results suggest that both chemogenetic LC stimulation and foot shock lead to similar changes in vmPFC activity.

### Locus coeruleus global chemogenetic stimulation impairs extinction learning

Studies exploring the effects of LC manipulations during extinction found conflicting results^14,27,30^. Therefore, we aimed to determine, under our parameters, how chemogenetic LC stimulation would affect delayed extinction. Animals from the previous experiment were re-conditioned and then underwent an extinction procedure after either VEH or CNO injections as shown in Figure 3B (groups were balanced to assure comparable freezing levels at the end of conditioning). During the extinction session, no significant differences in freezing behavior were observed between groups, but there was a trend towards higher freezing levels in CNO-treated animals (*F*_1,10_ = 3.86, *p* = 0.08). On the next day, animals were returned to the chambers for an extinction retrieval session. During retrieval, freezing was higher in CNO-treated animals, as confirmed by a *t*-test (*t* = 5.73, *p* < 0.01) (Figure 3C). These results suggest that chemogenetic stimulation of the LC leads to extinction impairment.

**Figure 3.**
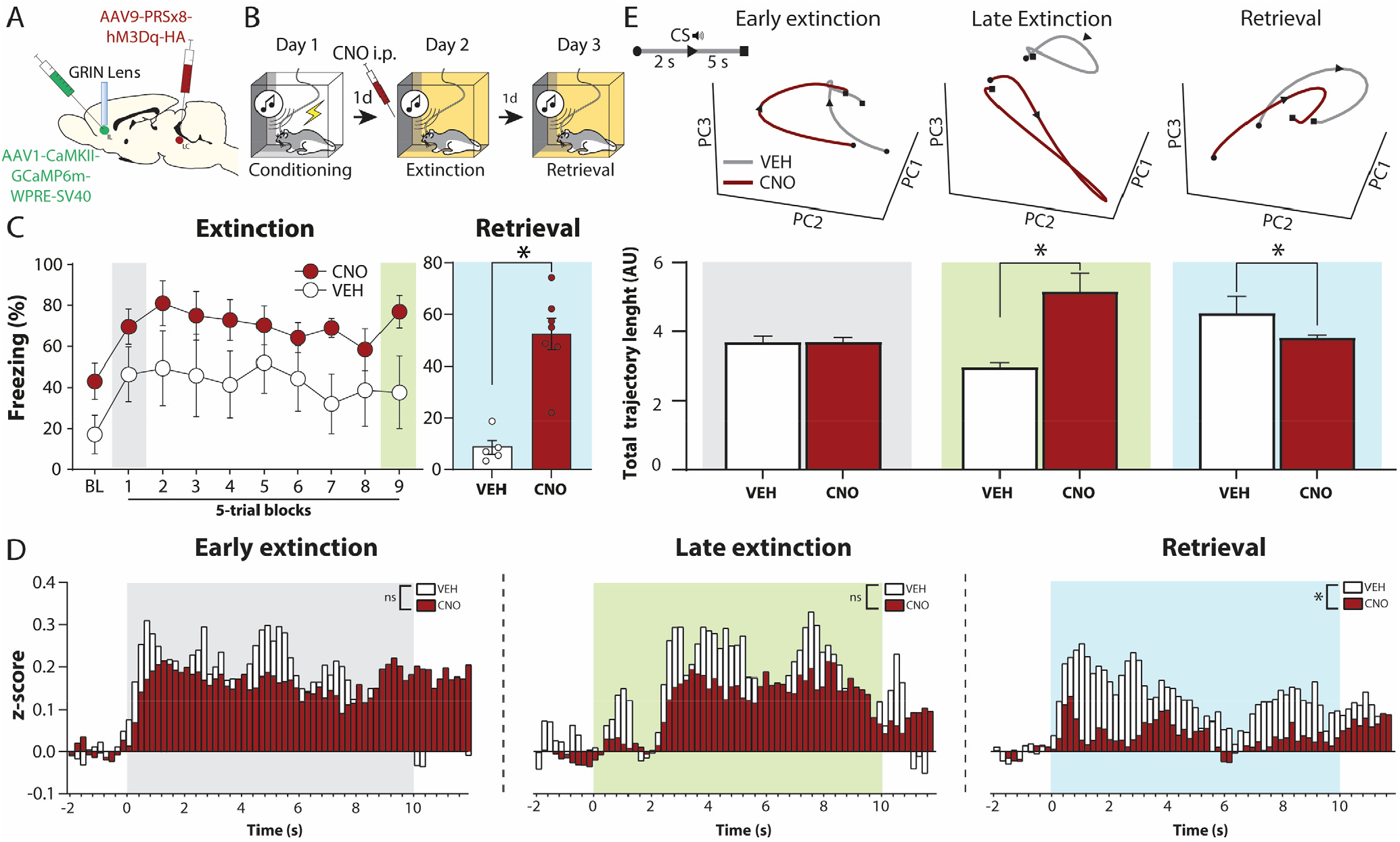
Chemogenetic activation of locus coeruleus impairs extinction learning. (A) Surgery schematic. A DREADD-carrying vector with specificity to NE+ neurons was injected in the LC, a GCaMP vector was injected in the vmPFC, and a GRIN lens was implanted on the vmPFC. (B) Behavioral schedule – animals were conditioned, and on the next day received injections of CNO (*n* = 7; 3 female and 4 male) or VEH (*n* = 5; 3 female and 2 male) 15 min before extinction and were tested on the next day in an extinction retrieval session. (C) The two groups did not have significant freezing differences during extinction, but CNO-treated animals had higher freezing during the retrieval session. (D) Peri-event time histograms for early extinction, late extinction and retrieval trials. During early extinction there were no significant differences in the average activity within the PETH window, but VEH-treated animals had higher average activity during late extinction and retrieval trials. (E) PCA was used to examine population activity during early extinction, late extinction and retrieval trials. The total trajectory length for VEH and CNO animals was not different for early extinction trials, but during late extinction CNO had a longer trajectory and during retrieval VEH animals had a longer trajectory. * = *p* < 0.05, CNO = clozapine-*N*-oxide, CS = conditioned stimulus, LC = Locus coeruleus, ns = not significant. PC = principal component, VEH = vehicle, vmPFC = ventromedial prefrontal cortex.

The first block of extinction is thought to correspond to a higher fear state, while late extinction and retrieval trials correspond to a lower fear state. Thus, we compared peri-event time histogram (PETH) for high and low fear state portions of extinction and retrieval from VEH and CNO-treated animals. To achieve that, we obtained the area under curve (AUC) for each animal and condition and then compared the average AUC from the two drug conditions. Although the groups did not differ at the beginning of extinction (*t* = 1.48, *p* = 0.20) and late extinction (*t* = 1.87, *p* = 0.12), retrieval trials had increased responses (*t* = 5.25, *p* < 0.01) in VEH-treated animals (Figure 3D).

To compare vmPFC population dynamics, we performed a principal component analysis (PCA) in all neurons to reduce dimensionality and visualize neural trajectories^31^.When neural trajectories during these trials were compared across groups, we observed no differences in total trajectory length during early extinction trials (*t* = 0.08, *p* = 0.94), while CNO led to higher trajectory lengths during late extinction trials (*t* = 6.58, *p* <0.01) and shorter trajectories during retrieval (*t* = 2.91, *p* = 0.03). Additionally, when early extinction and retrieval trajectory lengths are compared, there is no difference for the CNO condition (*t* = 1.44, *p* = 0.20), but there is a marginal effect in the VEH condition (*t* = 2.75, *p* = 0.05). These results suggest that LC activation affects how extinction is encoded in the vmPFC, which corresponds to lower CS-evoked responses during both late extinction and retrieval, and asymmetrical changes in population dynamics during late extinction trials and shorter trajectory during retrieval.

### Basolateral amygdala β-receptors are necessary for prefrontal modulation by locus coeruleus

Our results reveal that chemogenetic stimulation of the LC promotes freezing and suppresses the vmPFC. These results suggest that LC stimulation might act through the BLA to perturb vmPFC activity. To investigate this possibility, we implanted 4 rats with GRIN lenses in the vmPFC, a cannula in the BLA (ipsilateral to GRIN lens), and injected AAV9-FLEX-GCaMP8f in the vmPFC and AAV9-PRSx8-hM3Dq-HA in the LC (Figure 4A). Four weeks after surgery, animals underwent a within-subjects procedure to measure the effects of LC stimulation and intra-BLA propranolol (PROP): animals received either PROP or VEH intra-BLA injections, followed 5 min later by systemic VEH or CNO injections, and were tested 15 min after that (Figure 4B).

**Figure 4.**
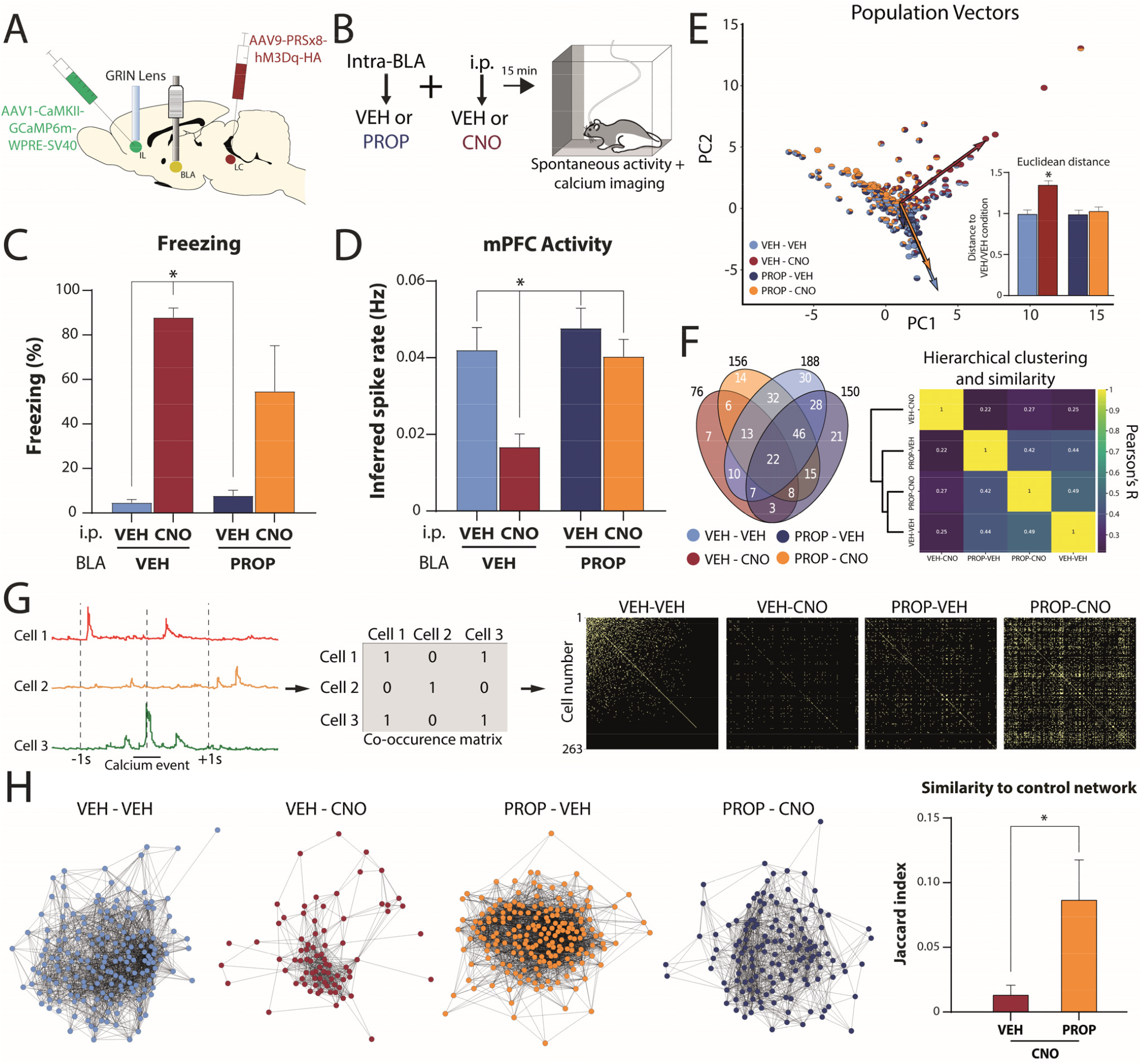
Locus coeruleus norepinephrine release modulates prefrontal activity via basolateral amygdala β-receptors. (A) Surgery schematic. A DREADD-carrying vector with specificity to NE+ neurons was injected in the LC, a GCaMP vector was injected in the vmPFC, a cannula was implanted unilaterally on the BLA, and a GRIN lens was implanted on the vmPFC, ipsilateral to the cannula (*n* = 4; 1 female and 3 male). (B) Behavioral procedure: animals received intra-BLA injections of PROP or VEH, followed by i.p. injections of CNO or VEH, and were placed in a chamber 15 min later for behavioral assessment and calcium imaging. Importantly, all animals were tested under all conditions in a within-subjects design. (C) Freezing behavior during the session. Both groups treated with CNO presented increased freezing levels, however, only the VEH x CNO group was significantly different from animals which did not receive CNO. (D) When the inferred spike rate was analyzed, we observed a marked decrease in vmPFC inferred spikes in the VEH-CNO group in comparison with all others. (E) Population vectors showing a scatter plot of all detected neurons in PCA space, with the population vector of each condition plotted on top of the individual neurons. Inset, averaged Euclidean distance of each neuron detected for each condition from the mean coordinates of the VEH x VEH group shows that the VEH x CNO group is significantly higher than all other groups. (F) Venn diagram and hierarchical clustering and similarity heatmap based on the detection of cells across conditions (VEH x VEH = 188 cells, VEH x CNO = 76 cells, PROP x VEH = 150 cells, PROP x CNO = 156 cells). (G) A co-occurrence matrix was generated for each condition based on neurons that had calcium events detected within the same 1 s windows, and each co-occurrence matrix was used to plot a network for every animal in each condition. (H) Network comparisons showing that the Jaccard similarity to the VEH x VEH network is reduced in VEH x CNO and rescued by propranolol and the clustering index for each network. BLA = basolateral amygdala, CNO = clozapine-n-oxide, i.p. = intraperitoneal, PC = principal component, PROP = propranolol, VEH = vehicle, vmPFC = ventromedial prefrontal cortex, * = *p* < 0.05, line and bar plots represent mean ± s.e.m.s.

As shown in Figure 4C, we observed a pronounced increase in freezing behavior in animals receiving CNO, regardless of the BLA treatment. This was confirmed by a two-way ANOVA, which found a main effect of CNO (*F*_1, 12_ = 15.39, *p* < 0.01) without interaction (*F*_1, 12_ = 2.34, *p* = 0.15). The Tukey’s *post hoc* revealed that the only significant pairwise differences found were between the VEHxCNO group and the VEHxVEH and VEHxPROP groups (*p*’s < 0.01). When average inferred spike rate was compared, we observed that the CNOxVEH group had lower inferred spikes than all others (Figure 4D). This was confirmed by a two-way ANOVA that showed a significant main effect of CNO (*F*_1, 53_ = 13.73, *p* < 0.01) and of PROP (*F*_1, 53_ = 10.92, *p* < 0.01), and a trend towards a significant interaction between CNO and PROP (*F*_1, 53_ = 2.88, *p* = 0.10). The Tukey’s *post-hoc* test that showed a significant difference between the CNOxVEH group and all others (*p’s <* 0.01), while the three other groups were not different among themselves (*p’s* > 0.91). Additionally, we have found an increase in BLA NE signaling followed by footshock, that was blunted by systemically injected clonidine (Figure S2), as the peak response to footshock was smaller in clonidine-treated animals (*t* = 4.63, *p <* 0.01). Taken together, these data show that NE release in the BLA is increased by footshock stress as well as LC stimulation, and BLA NE receptor activation is blunted by a α2 agonist, and the reduction in vmPFC activity induced by LC stimulation requires β-receptor activation.

Additionally, we performed a PCA to further compare vmPFC dynamics at the population level and visualize vmPFC ensembles in a low-dimensional space. We used all neurons detected during at least one of the four sessions across all animals and performed the PCA on the entire dataset. When the population vector for each condition was plotted in a two-dimensional space using the first two PCs, the CNOxVEH vector was orthogonal to the other vectors (Figure 4E). To statistically compare the differences in population dynamics, we measured the Euclidean distance to the mean coordinates of the VEHxVEH condition for each cell and found that the VEHxCNO had a higher distance from the VEHxVEH condition than all the other groups (Figure 4E, *inset*). That was confirmed by a two-way ANOVA which revealed a significant effect of CNO (*F*_1,564_ = 7.12, *p* < 0.01), PROP (*F*_1,564_ = 4.80, *p* = 0.03), with a significant interaction between the factors (*F*_1,564_ = 4.59, *p* = 0.03). Additionally, we generated a Venn diagram to visualize the number of neurons detected during each condition (Figure 4F, *left panel*), then a similarity matrix and hierarchical clustering of conditions was generated based on the detection of the same cells across days (Figure 4F, right panel). Hierarchical clustering showed that three conditions (VEHxVEH, PROPxVEH, and PROPxCNO) clustered together, while VEHxCNO was isolated in a separate branch, suggesting a distinct pattern of neuronal activation compared to the other three conditions.

To investigate the effects of LC activation and intra-BLA propranolol on vmPFC networks, we extracted calcium events of individual neurons across time and created a co-occurrence matrix based on the co-activation of neuron pairs within 1-s windows. For each animal, one network was generated per condition (Figure 4G). We then measured the Jaccard similarity index between the VEHxCNO and PROPxCNO network and the VEHxVEH network for each. We found that the vmPFC networks for the PROPxCNO condition were significantly more similar to the VEHxVEH condition when compared to the VEHxCNO condition (Figure 4H). That was confirmed by a t-test (t = 2.66, p = 0.04). When networks were analyzed in isolation using the clustering coefficient (Figure S3), we found that the VEHxCNO condition had networks with lower clustering coefficients when compared to the VEHxVEH networks (t = 14.41, p < 0.01), while the PROPxVEH and PROPxCNO did not have significant differences in the clustering coefficients (*t* = 0.59, *p* = 0.61).

Together, our data suggest that LC activation-induced freezing is partially rescued by β-receptor blockade in the BLA, and inferred firing rate is fully restored to control conditions. Similarly, population dynamics and vmPFC network activity are also dysregulated by LC stimulation and restored by intra-BLA propranolol injections.

### LC chemogenetic stimulation increases activity in vmPFC-projecting BLA neurons

The previous experiments suggest that the effects of LC stimulation on freezing behavior and extinction involve the BLA. To further explore that possibility, thirteen rats received injections of AAV9-PRSx8-hM3Dq-HA or AAV9-hSyn-GFP in the LC, AAV9-FLEX-GCaMP8f in the BLA, and CAV2-Cre in the vmPFC, so that we could record from vmPFC-projecting BLA neurons while the LC was chemogenetically activated (Figure 5A). In this experiment, we combined LC stimulation with a weak fear conditioning protocol (Figure 5B). This protocol has been used by our lab in the past and does not lead to IED by itself but does so when combined with LC stimulation^14^.

**Figure 5.**
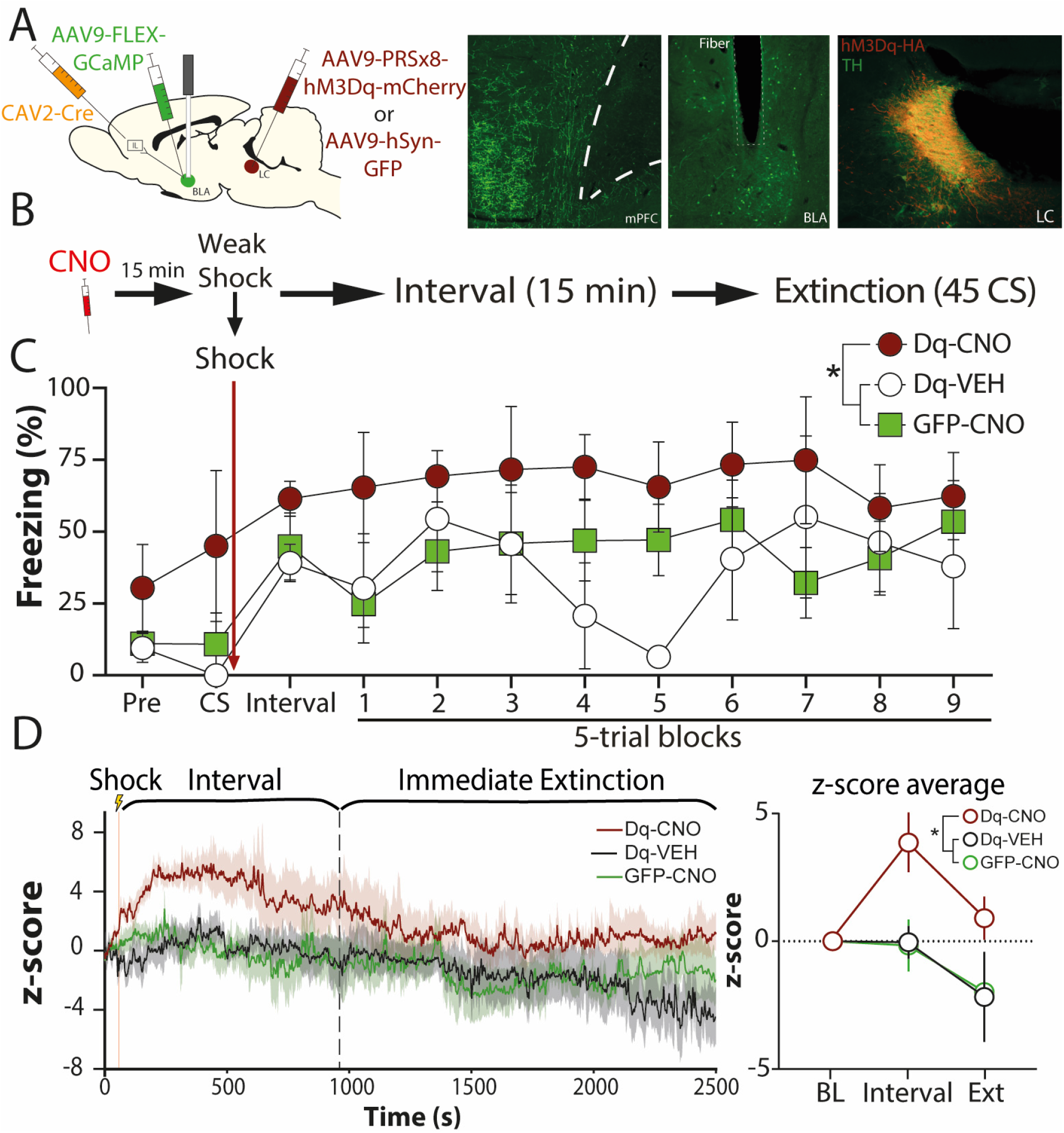
vmPFC-projecting BLA neurons increase activity during immediate extinction in response to locus coeruleus stimulation. (A) Surgery schematic and representative histology. A DREADD-carrying vector with specificity to NE+ neurons, or a control vector was injected in the LC, a retrograde Cre vector was injected in the vmPFC, a Cre-dependent vector was injected into the BLA, and an optic fiber was implanted on the BLA for fiber photometry recordings. (B) Behavioral schedule – six weeks after surgeries, animals received injections of CNO or VEH and underwent a weak conditioning protocol followed by an immediate extinction session 15 min later. (C) During the entire session, the Dq-CNO group (*n* = 4; 2 females and 2 males), which had the active DREADD and received the CNO treatment had higher freezing levels when compared to the other groups, which controlled for CNO (GFP-CNO; *n* = 3; 2 males and 1 female) and virus infection in the LC (Dq-VEH; *n* = 6; 4 females and 2 males). (D) Fiber photometry data obtained across the entire session (l*eft panel)* shows that the Dq-CNO (*n* = 6 channels) group had higher calcium transients during the post-shock interval and the extinction session (*right panel*) when compared to Dq-VEH (*n* = 5 channels) and GFP-CNO (*n* = 6 channels). BL = baseline, BLA = basolateral amygdala, CNO = clozapine-*N*-oxide, VEH = vehicle, * = *p* < 0.05, line and bar plots represent mean ± s.e.m.s.

Six weeks after surgery, animals received i.p. injections of VEH or CNO 15 min prior to fear conditioning and remained in the cage for another 15 min until extinction started. Fiber photometry data were acquired throughout the session. As shown in Figure 5C, animals expressing hM3Dq vector in LC and injected with CNO had higher freezing throughout the session. When freezing data for the entire session was averaged, a main effect of group was revealed by an one-way ANOVA (*F*_2, 33_ = 12.46, *p* < 0.01). Tukey’s *post hoc* comparisons showed that although the Dq-VEH and GFP-CNO groups were not different from each other (p = 0.65), both were significantly different from the Dq-CNO group (p’s < 0.01).

To examine calcium transients during the entire session, we used the averaged z-score for each stage of the session (BL, post-conditioning interval, extinction) (Figure 5D). A two-way ANOVA performed on these data revealed a significant main effect of group (F_2, 12_ = 4.62, p = 0.03) and stage of the session (F_2, 24_ = 6.60, p < 0.01), without an interaction between the factors (F_4, 24_ = 2.35, p = 0.08). A Tukey’s post hoc comparison showed that although no groups were different during the baseline period (p’s > 0.99), the Dq-CNO group had higher z-scores than controls during the post-conditioning interval (p’s < 0.01). During extinction calcium activity in this group was significantly higher than that in the Dq-VEH group (p = 0.049) and had a trend towards significance when compared to the GFP-CNO group (p = 0.07).

These results suggest that although a weak shock per se is not sufficient to increase activity in vmPFC-projecting BLA neurons, it drives higher activity in those neurons when combined with LC stimulation. This might be the mechanism through which LC chemogenetic stimulation causes the IED in a paradigm that does not lead to it.

### Activation of LC ⟶ BLA projections is sufficient to drive the immediate extinction deficit

To determine whether activation of LC⟶BLA projections in isolation impairs extinction learning and retrieval, we used another set of rats to express hM3Dq specifically in BLA-projecting LC neurons. Thirty rats received intra-BLA injections of the retrograde viruses CAV2-PRSx8-hM3Dq-mCherry or CAV2-mCherry (Figure 6A). Six weeks later, animals underwent a weak fear conditioning protocol under VEH or CNO followed by an immediate extinction procedure as described above (Figure 6B). As shown in Figure 6C, there was not a significant increase in freezing in the post-shock period in animals receiving VEH or blank virus. However, CNO significantly increased freezing in animals expressing hM3Dq. That was confirmed by a two-way ANOVA that revealed a main effect of virus (F_1, 52_ = 16.63, p < 0.01), CNO (F_3, 52_ = 4.16, p < 0.01), with a significant interaction between the factors (F_3, 52_ = 3.70, p = 0.02). Shortly after conditioning (15 min), animals underwent an extinction session. No differences were observed during baseline (F_3, 26_ = 1.66, p = 0.20). When the entire extinction session was analyzed, there was a main effect of trial (F_9, 234_ = 3.47, p < 0.01), but no other significant main effects or interactions were found (F’s < 1.13, p’s > 0.35). However, when the first two blocks were analyzed in isolation, a two-way ANOVA revealed a significant interaction between virus and drug (F_1, 26_ = 6.86, p = 0.01), suggesting that CNO increased freezing only in hM3Dq animals. Two days later, the animals had a retrieval test, in which the hM3Dq-CNO group froze significantly more than the other groups, as confirmed by a significant interaction between virus and drug (F_1, 26_ = 4.76, *p* = 0. 04). These results show that LC⟶BLA stimulation acutely increases freezing specifically after footshock and tone presentation in the immediate extinction session, but not during extinction baseline, which suggests that the activation of this pathway is increasing fear responses to threatening stimuli, but not generally increasing freezing like observed for LC global stimulation in the first two experiments. Additionally, as differences persisted during retrieval, these results suggest that LC⟶BLA stimulation promotes the IED.

**Figure 6.**
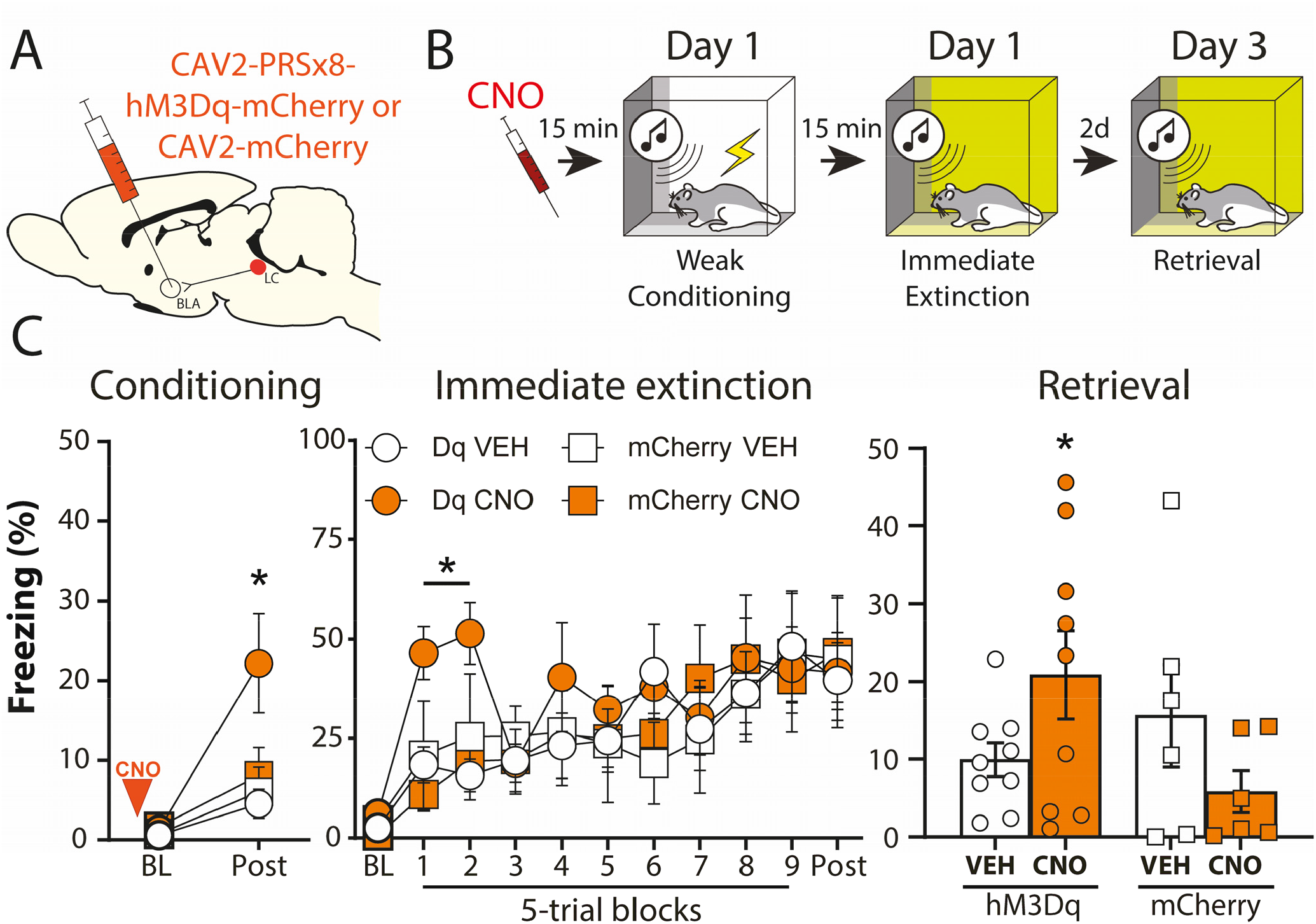
Stimulation of BLA-projecting LC neurons promotes the immediate extinction deficit. (A) Surgery schematic. (B) Behavioral schedule six weeks after surgeries, animals received injections of VEH or CNO, and underwent the same immediate extinction procedure as described above in panel B. (C) The animals that received injections of hM3Dq-carrying viruses and CNO injections had higher freezing levels during conditioning, immediate extinction and retrieval when compared to the other groups (*n*’s per group: hM3Dq-VEH = 9; 5 females and 4 males; hM3Dq-CNO = 9; 4 females and 5 males; mCherry-VEH = 6; 3 females and 3 males; mCherry-CNO = 6; 3 females and 3 males). BL = baseline, CNO = clozapine-n-oxide, VEH = vehicle, * = *p* < 0.05, line and bar plots represent mean ± s.e.m.s.

## DISCUSSION

Here we show that LC chemogenetic stimulation increases freezing behavior and reduces neuronal activity in the vmPFC. The effects of LC stimulation on vmPFC activity resemble those of footshock, commonly used as the unconditioned stimulus in fear conditioning. These results suggest that LC activation acts as a stress signal and shifts brain wide activity and behavior to a heightened arousal state. We then explored how the projections from the LC to the BLA are involved in these effects and in extinction learning. We found that the activation of the LC impairs delayed extinction and activation of LC⟶BLA projections promotes the IED. Additionally, blocking BLA β-receptors prevents disruption of vmPFC activity and population dynamics caused by LC stimulation. Finally, we show that LC activation during a weak fear conditioning procedure drives increased activity in BLA⟶vmPFC projection neurons, which are implicated in the immediate extinction deficit^32^. Together, our results suggest that footshock and LC activation drives NE release in the BLA and increases activity in vmPFC-projecting BLA neurons to disrupt prefrontal activity and generate a high stress state, which in turn leads to extinction deficits.

Interestingly, we found that LC stimulation not only reduces vmPFC spike firing but also affects population and network dynamics in the vmPFC. When we compared the proportions of neurons modulated by footshocks and chemogenetic LC activation, we found that most cells reduced their activity in response to both CNO and footshock. Additionally, when we blocked BLA β-receptors in combination with LC stimulation, we found that the latter shifted population dynamics and dysregulated vmPFC networks and blocking BLA β-receptors was able to rescue those phenotypes. When population dynamics were assessed throughout extinction, differences emerged at the end of extinction and persisted to the retrieval session. The neural trajectory length over time can be used as an index of neural dynamics^33^. Our results therefore suggest that LC activation led to increased dynamics at the end of extinction and lower dynamics during retrieval, even though we observed a reduction in vmPFC spike firing and higher freezing levels at both time points.

Several studies have suggested an inverse relationship between stress and the success of extinction learning^13^. A common neurobiological substrate that emerges after acute or chronic stress models is reduced vmPFC activity^10,34–38^. The LC-NE system is involved in stress processing, as well as in extinction learning^10,14–16,27^ Considering the importance of NE release across the brain for cognitive processes and the role of the LC in adjusting brain function in response to sensory stimulation and interoceptive cues, the LC is perfectly positioned to be a conduit between stress and extinction deficits.

In opposition to our findings, others have shown that global optogenetic inhibition of the LC impairs extinction retrieval without affecting behavior acutely^39^. In our case, chemogenetic stimulation led to a drastic acute effect on freezing and impaired extinction retrieval on the next day when the animals were tested in a drug-free state. Interestingly, that study revealed extinction- and fear-responsive LC ensembles, and a dissociation between LC⟶BLA and LC⟶vmPFC projections, where inactivation of the BLA projection promotes extinction and inactivation of LC⟶vmPFC projections impairs it^27^. This is in line with our observations showing that chemogenetic stimulation of the LC⟶BLA pathway impairs extinction learning and shown that LC activation suppresses vmPFC activity through the BLA. Different LC projections have different effects on freezing and extinction, and the different techniques used (i.e. optogenetics and chemogenetics) could be differentially affecting these efferents.

It has been suggested that NE is necessary for extinction^40–45^. From this perspective, the finding that chemogenetic activation of the LC impairs extinction may be surprising. One possibility is that although NE is necessary for memory acquisition and consolidation, the effects of LC activation on learning might reflect an inverted-U shaped curve. An inverted U-shaped curve between arousal and performance has been reported many times and is often referred to as the Yerkes-Dodson law, and of course NE is tightly related to arousal regulation. Importantly, the integrative model of LC function proposed by Aston-Jones & Cohen (2005) assumes that LC activity and cognitive performance also have an inverted-U shaped curve dynamic. Additionally, there are also reports of inverted-U shaped curves for NE and fear learning^46,47^. Therefore, our LC stimulation parameters may shift the system to a high-tonic level of activity which increases anxiety and stress and reduces attention to the contingency change that happens during extinction learning, and therefore extinction fails to happen.

It is also important to note that in our experiments involving immediate extinction, we are performing manipulations within the consolidation time window of the recently formed fear memory. Indeed, there are numerous reports showing that BLA NE affects fear memory consolidation^48–53^. On the other hand, there are also reports, including studies from our laboratory, showing that β-receptor blockade or activation in the BLA do not affect fear memory consolidation^15,54–56^. Unfortunately, this is an unavoidable confounding factor when studying the IED, which is indissociable from the high-stress levels that linger after conditioning^57^. At any rate, the bulk of evidence suggests that LC-NE manipulations do not affect memory encoding but rather involve global arousal and stress state transitions. This perspective is supported by the β-receptor blockade experiment, in which animals were never conditioned, but BLA β-receptors were found to affect freezing and vmPFC population activity and dynamics (Figure 4).

In sum, our findings demonstrate that LC stimulation induces a high-stress state characterized by vmPFC suppression, increased NE release in the BLA, and activation of the BLA⟶vmPFC pathway. Activation of this circuit increases freezing and impairs extinction learning. These effects parallel those observed in stress-related extinction deficits, suggesting that the LC serves as a critical mediator between stress and behavior. Furthermore, our findings support the hypothesis that elevated LC activity induces brain-wide alterations that increase arousal beyond optimal levels, impairing cognitive flexibility and extinction. Together, this work advances our understanding of how stress impacts extinction learning and underscores the importance of the LC-NE system and its targets, such as BLA β-receptors, as potential mechanisms for therapeutic intervention in stress-related disorders such as PTSD.

## ACKNOWLEDGEMENTS

We thank Morgana Favero and Waylin Yu for their technical assistance with the miniscope experiment, Elena Vazey for providing us viral vectors used in this study, and Flávio Afonso Gonçalves Mourão for his help with statistical analyses.

## AUTHOR CONTRIBUTIONS

AB, SS, HB, and SM designed the miniscope experiments; HB, VJ, AB and SM designed the fiber photometry and LC⟶BLA pathway experiments. HB, AB, and SS analyzed the data. AB, SS, and HB performed the miniscope experiments, HB, VJ, SP and LF performed the fiber photometry experiments, HB, VJ, and AB performed the LC⟶BLA pathway experiment. HB and SM wrote the manuscript.

## DATA AND CODE AVAILABILITY

The data from these experiments and codes used for analyses are available from the corresponding author upon request.

## SUPPLEMENTARY RESULTS

**Supplemental figure 1:**
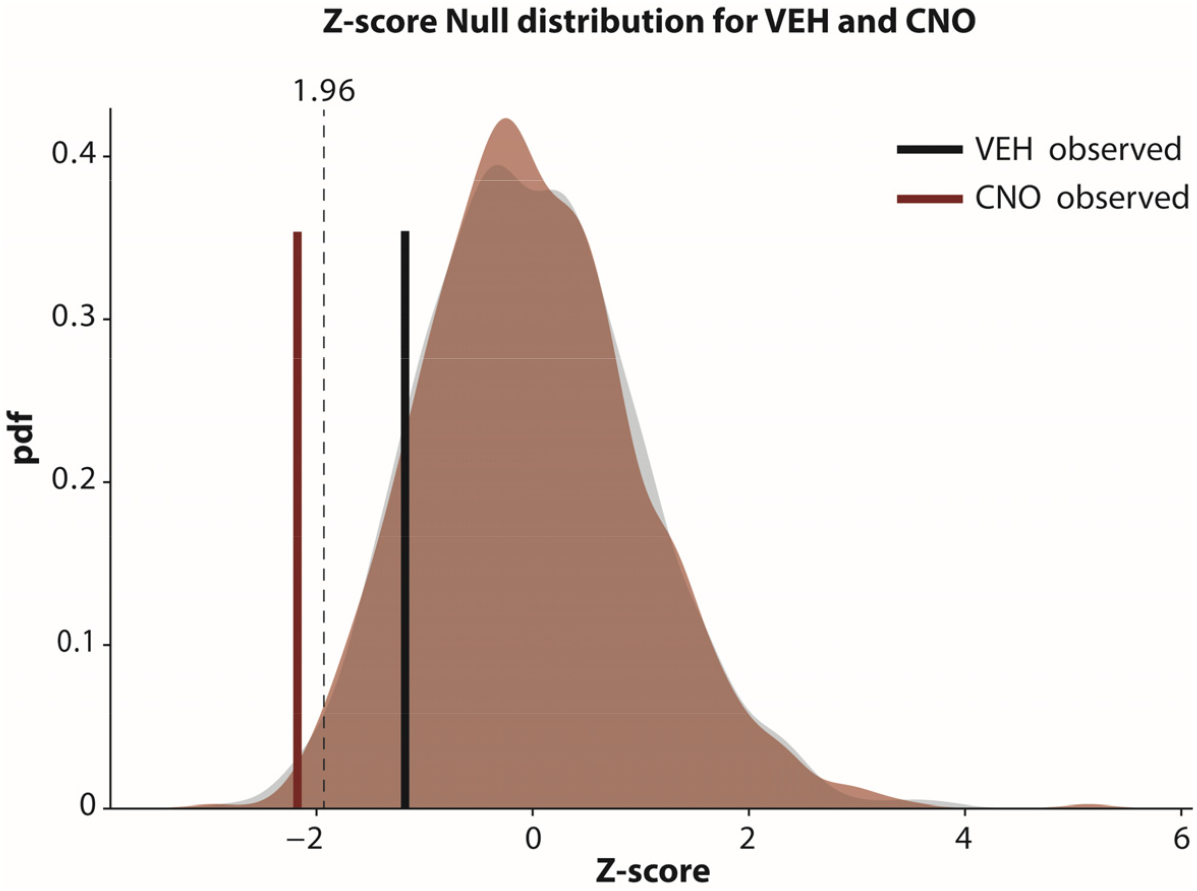
Permutation analysis of spontaneous inferred spike rate after VEH or CNO injections. (A) To assess whether observed changes in activity between baseline and post-treatment periods were statistically significant, we used a permutation test. For each condition (VEH and CNO), we computed the ratio of mean post-treatment activity to mean baseline activity for each cell. We then generated a null distribution of these ratios by randomly permuting baseline and post-treatment labels across the entire population 1,000 times and used these iterations to obtain new shuffled ratios. The observed ratio was transformed into a Z-score based on the mean and standard deviation of the null distribution, and a p-value was calculated from the standard normal distribution corresponding to a two-tailed test. We found that while the VEH observed ratio did not have a significant difference from the null model, the CNO observed ratio was significantly lower than the null model.

**Supplemental figure 2:**
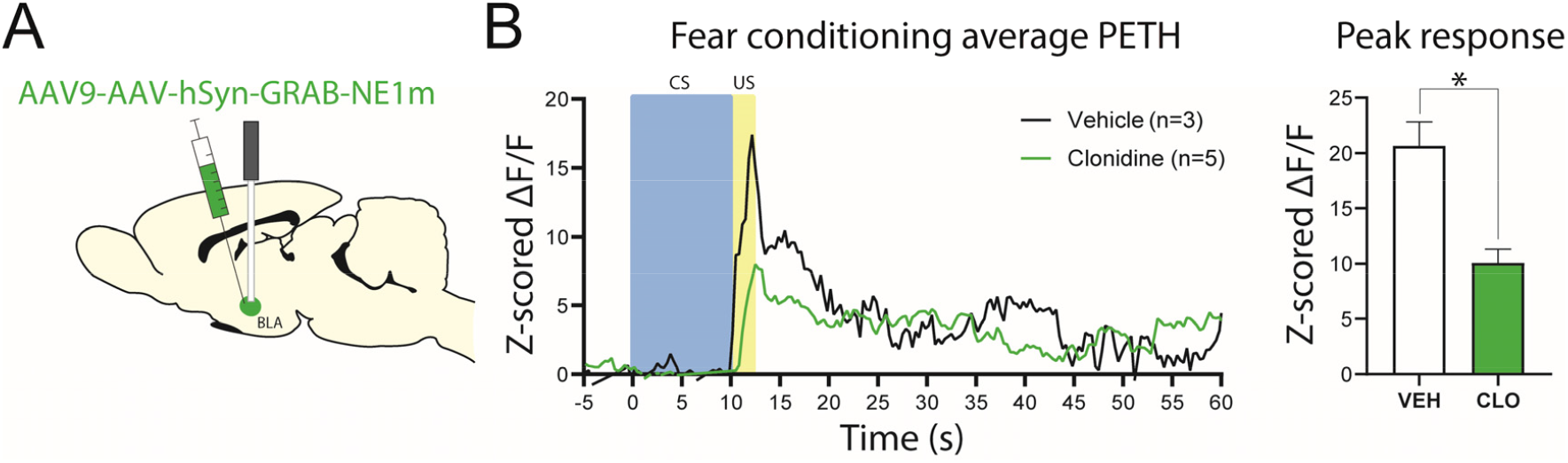
Fear conditioning leads to an increase in norepinephrine release in the basolateral amygdala. (A) Surgery schematic. (B) Four weeks after surgeries, animals received clonidine and underwent a fear conditioning session 20 minutes later. Fiber photometric data were normalized and ΔF/F signals were z-scored based on the 15 seconds preceding shock onset. (C) The peak response for each animal was extracted from the Z-scored trace and those were averaged for each group. * = p <0.05, BLA = basolateral amygdala, CLO = clonidine, CS = conditioned stimulus, US = unconditioned stimulus, VEH = vehicle.

**Supplemental figure 3:**
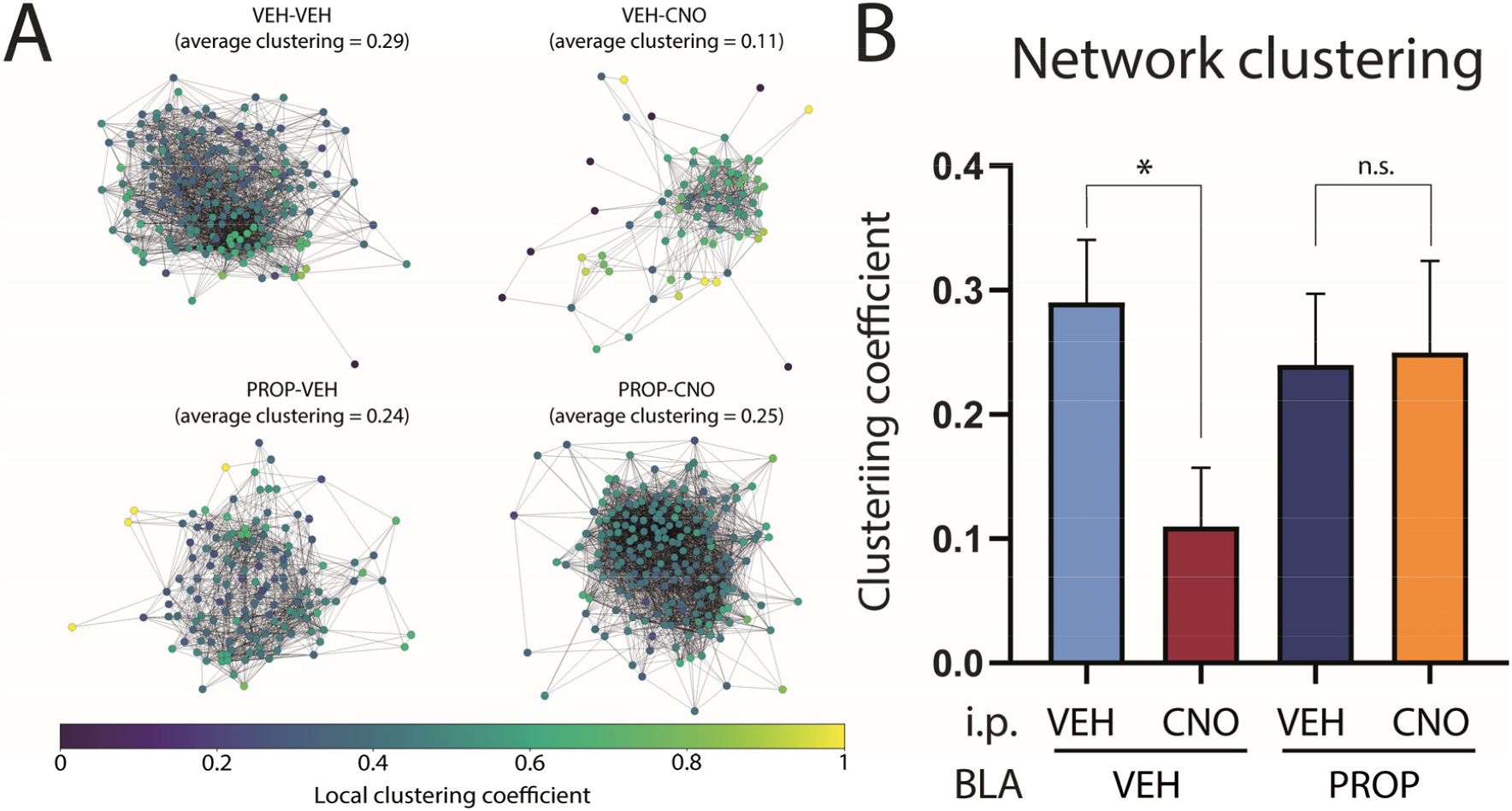
Locus coeruleus chemogenetic stimulation disrupts vmPFC network clustering. (A) Representative networks for each group showing the local clustering coefficients for individual nodes. (B) The individual clustering coefficients were averaged for each network to obtain a global clustering coefficient for each condition. While the VEH-CNO condition had a significantly lower clustering coefficient when compared to the VEH-VEH condition, that was not found when PROP-treated VEH and CNO animals were compared. * = p <0.05, BLA = basolateral amygdala, CNO = clozapine-n-oxide, i.p. = intraperitoneal, PROP = propranolol.

